# Transcriptome-Wide Analysis of HuR Function Identifies TXNIP-Mediated Redox Dysregulation in Osteocytes

**DOI:** 10.1101/2025.08.08.668956

**Authors:** Ziqiu Fan, Aseel Marahleh, Hideki Kitaura, Jiayi Ren, Abdulrahman Mousa, Fumitoshi Ohori, Kuniyasu Niizuma, Sherif Rashad, Hiroyasu Kanetaka

**Affiliations:** Orthodontics and Dentofacial Orthopedics Department, Graduate School of Dentistry, Tohoku University, Japan; Frontier Research Institute for Interdisciplinary Sciences, Tohoku University, Japan; Center for Environmental Response and Aging, Institute of Development, Aging and Cancer, Tohoku University.; Department of Neurosurgical Engineering, Graduate School of Biomedical Engineering, Tohoku University; Department of Translational Neuroscience, Graduate School of Medicine, Tohoku University

**Keywords:** HuR/Elavl1, RNA binding proteins, alternative splicing, hyperglycemia, osteocytes, TXNIP, translation, mTOR

## Abstract

Post-transcriptional gene regulation is central to maintaining cellular homeostasis, among its mechanism alternative splicing (AS) fine-tunes cellular adaptation to stress. In this study we employed an approach combining RNA splicing analysis to define RNA binding protein (RBP) motif enrichment around alternatively spliced exons in primary osteocytes cultured in high glucose conditions (HG). We identified the RBP human antigen R (HuR) as a top candidate regulator of AS. Loss of HuR reshaped the transcriptome through widespread changes in gene expression and splicing, converging on two major pathways: stress response and translational control. Functional validation of splicing revealed that HuR depletion heightened oxidative stress sensitivity and compromised cell viability under HG by stabilizing and upregulating TXNIP, a thioredoxin inhibitor. HuR knockdown also impaired mitochondrial mass and function and disrupted key translational signals, despite preserving global protein output. These findings establish HuR as a central post-transcriptional regulator of osteocyte survival and metabolic adaptation under high glucose stress, with potential implications for hyperglycemic bone fragility.

## Introduction

Post-transcriptional gene regulation modulates gene expression by modulating RNA processing through RNA splicing, modifications, localization, stability, translation and fat,^1,2,3,4,5,6^ generating transcript variants that contribute to proteomic diversity. Thus, post-transcriptional regulation plays a key role in stress adaptation optimizing protein synthesis to promote cell survival.^7,8^

RNA binding proteins (RBPs) regulate RNAs by binding structured RNA motifs to control nearly every aspect of RNA metabolism and function.^9,10,11,12^ RBPs contribute to tissue-specific disease phenotypes, often reflecting the expression pattern of their RNA targets and cofactors as well as the RBP’s own abundance and activity level.^13^ Alterations in RBPs or their targets are associated with diverse pathologies, including neurodegenerative, muscular, autoimmune, and metabolic diseases driven by impaired RNA transport, splicing, or clearance, highlighting the critical role of RNA-RBPs interactome in determining disease outcomes.^9,13,14^

HuR [‘human antigen R’, also known as ELAVL1 (embryonic lethal abnormal vision-like 1)] is a ubiquitously expressed RNA-binding protein, a classic RBP, possessing RNA recognition motifs (RRM) and a key modulator of post transcriptional gene regulation.^15^ It has been extensively studied as a cancer drug target ^16^ and its global deletion has been shown to be embryonically lethal,^17^ and conditional knockouts experienced impaired post-natal formation of skeletal, immune, and other tissues.^18,19,20,21^ Beyond development, tissue-specific knockdowns disrupt metabolic and immune homeostasis; where glucose and lipid metabolism linking it to obesity and insulin resistance.^22,23,24^

In bone, an ovariectomy-induced bone loss model, had reduced HuR expression, whereas overexpression alleviated bone loss and silencing impaired osteoblast differentiation.^25^ In contrast, diabetic mouse models showed increased HuR mRNA levels, and its knockdown improved glucose metabolism and bone microarchitecture.^26^ Similarly, in MC3T3-E1 cells exposed to high glucose, HuR expression increased, and its silencing reduced apoptosis while promoting osteogenic differentiation.^26^ These studies demonstrate that HuR expression and its effects are tissue and context dependent.

Osteocytes, comprising the majority of bone cells, are post mitotic mechanosensors maintaining bone remodeling through orchestrating resorption and formation. They are endocrine and signaling cells that maintain skeletal homeostasis.^27,28,29,30^ To date, post-transcriptional RNA regulation in osteocytes and the RBPs regulating it remain largely uncharted. Here, we performed an unbiased RBP motif enrichment analysis on differentially spliced events from RNA-seq data of primary osteocytes from Dmp1-topaz mice cultured under high-glucose conditions. We introduce for the first time the alternative splicing programs regulated in osteocytes and map the RBPs that regulate them. We further reveal HuR as a key regulator during high glucose (HG) stress. To define its functional role, we employed an HuR knockdown model and investigated both differential gene expression and alternative splicing programs, thereby homing in on HuR’s role in sustaining cellular homeostasis, coordinating translation, and supporting mitochondrial function. Our data further demonstrates that HuR acts on thioredoxin-interacting protein (TXNIP) expression, a glucose-responsive inhibitor of the antioxidant thioredoxin and its silencing protect against oxidative stress injury.^31^ Specifically, HuR directly binds TXNIP (mRNA) and regulates its stability, thereby preserving redox balance. Collectively, these findings provide mechanistic insight into HuR’s function and highlight its potential role in preserving skeletal homeostasis in metabolic diseases.

## Results

### Splicing-driven RBP motif analysis identifies HuR as a top regulator in osteocytes under high glucose stress

Primary osteocytes obtained from Dmp1-Topaz mice were cultured in HG or NG for 72 hours. RNA-seq analysis showed minimal gene expression changes with TXNIP and CD36 among the top regulated (Figure 1A). rMATs analysis of the 5 canonical splicing patterns; skipped exon (SE), mutually exclusive exons (MXE), retained introns (RI), alternative 5’ splice site (A5SS), and alternative 3’ splice site (A3SS) revealed significant changes (FDR < 0.05, |Ψ| > 0.2) in ∼2% of total events, predominantly SE and A3SS (Figure 1B).^32^ GO Biological Processes (GOBP) and pathway enrichment analysis of AS showed that different AS programs contribute differently to cellular biology. Specifically, SE DSGs revealed enrichment in cytoskeletal remodeling and protein regulation via ubiquitination, and several pathways involved in matrix maintenance and repair (Figure 1C, Supplementary Table 1). MXE DSGs were enriched in pathways involved in collagen synthesis and organization, skeletal tissue development and pathways involved in cell growth and regulation of fluid balance (Figure 1D, Supplementary Table 1). A5SS DSGs were predominated by pathways related to post transcriptional and translational regulation including ribosome targeting, rRNA processing and quality control of mRNA translation. A3SS DSGs were involved in RNA splicing, several signaling pathways involved in cell growth and differentiation and parathyroid hormone action. DSGs in RI showed cell cycle related terms, ribosome and translation regulation as well as ubiquitination (Supplementary Fig 1A-C and Supplementary Table 1).

**Figure 1:**
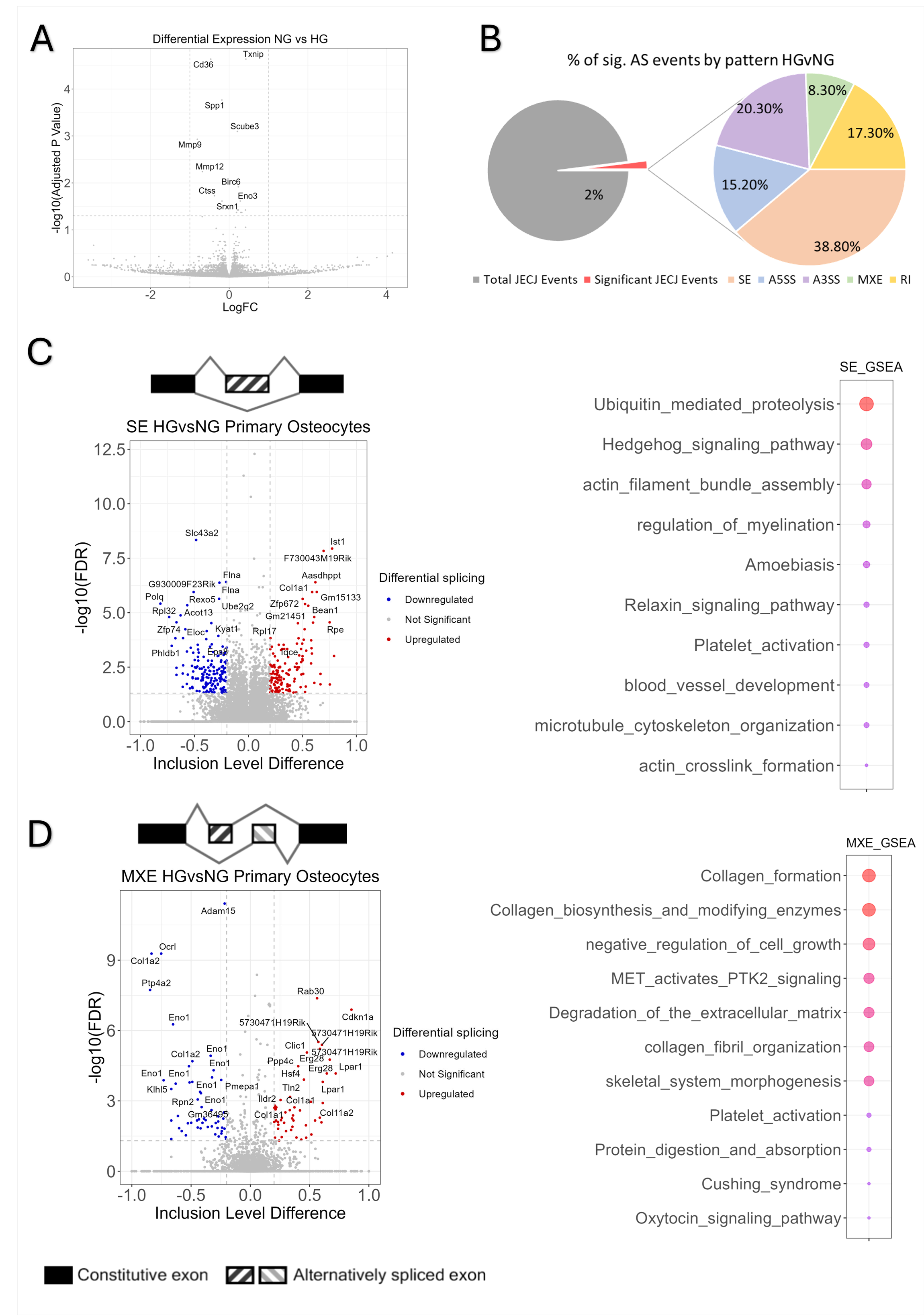
High glucose induces minimal differential gene expression and extensive AS changes in primary osteocytes. **(A)** The Volcano plot comparing the log2 FC and -log10 adjusted P value of RNA-seq data in primary osteocytes with genes showing the highest changes labelled. **(B)** The parent pie chart shows significant AS events (2%) of the total events at target and junction reads (JCEC) and the secondary pie chart shows the percentage of significant AS events of each type (SE, A5SS, A3SS, MXE, RI) from total significant events in primary osteocytes. **(C-G)** Volcano plots and overrepresentation analysis of DSGs around AS events showing inclusion level (Δ1) difference comparing 5.5mM and 25mM glucose at FDR < 0.05 and 0.2 < Δ1 < -0.2 in primary osteocytes. Overrepresentation analysis was done through the eVITTA web-based toolbox. All analysis was done on n=3. Also, see Supplementary Table 6.

Because these AS changes suggested altered post-transcriptional control, we next interrogated RBP motif enrichment using rMAPS2.^33^ We filtered the top up enriched and down enriched motif positions from all 5 AS datasets based on p values < 0.05 (t test). Around upregulated splice events, HNRNPL and HuR were the top enriched in SE (Figure 2A, Supplementary Table 2), in MXE, ZC3H14 and HuR. In A5SS, HNRNPL and HuR, in A3SS, FUS and HNRNPH2 and in RI we observed no enriched RBP motifs at a cut off value of p <0.05 (Supplementary Fig 2A-D and Supplementary Table 2). Around downregulated splice events, KHDRBS1 and ZC3H14 were the top enriched in SE (Figure 2B), in MXE, ZC3H14 and Tra2-beta, in A5SS, HNRPLL and HuR showed the highest enrichment. In A3SS, RBM47 and SRp20 were the most enriched while in RI RBMS3 and PABPN1 were dominant (Supplementary Fig 3A-D and Supplementary Table 2).

**Figure 2:**
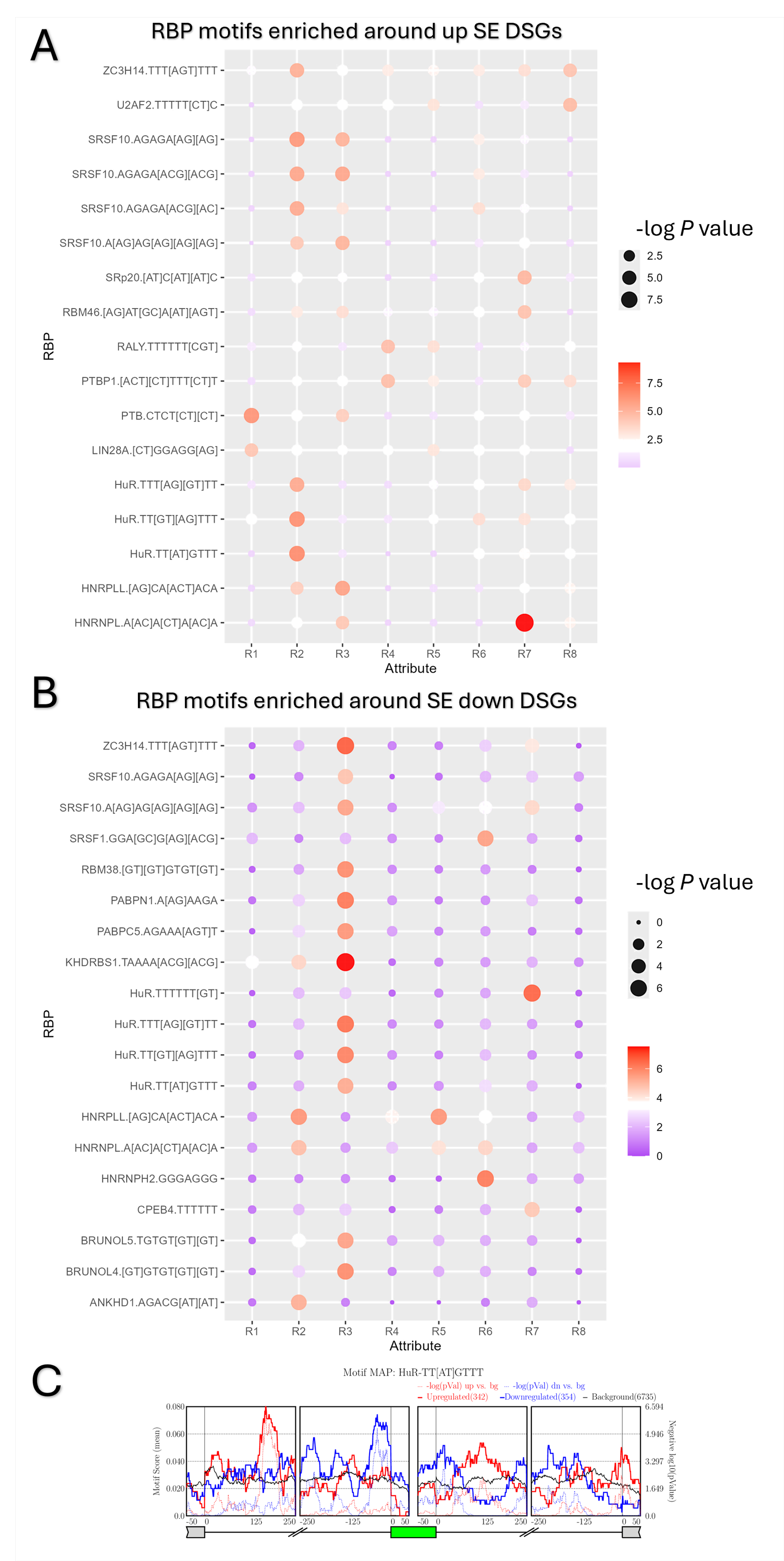
RBPs motifs significantly enriched around AS events. (A-B) The x axis (Attribute) represents the position (R) of each splicing event in 5’ → 3’. RBP binding motifs are described in the y axis. The panels show SE RBP motifs around up and down regulated DSGs and bubble size and color denote the – log10 P value. (C) Example positional distribution map of the HuR binding motif around target SE exon. Positional distribution maps were generated using rMAPs2. Intronic regions are 250-bp flanking sequences, and exon regions are 50-bp sequences from the start or end of the exon where the RBP binds around the spliced exon (green) and constitutive exons (grey). The solid red and blue lines are the motif enrichment scores, and the dotted lines are the p values in up and down enriched motifs, respectively. Black line refers to background RBPs distribution. Also, see Supplementary Table 7.

The analysis showed that several HuR motifs were enriched in upregulated splice events. Figure 2C shows an example of HuR binding motifs around SE differentially spliced exons. Our bioinformatics analysis points to HuR as a top regulatory RBP in AS, therefore we measured HuR mRNA and protein levels in MLO-Y4 cells exposed to HG. mRNA expression was downregulated as measured by qPCR while protein expression did not change as measured by western blot (Figure 3A-C). Similarly, HuR mRNA also decreased in the long bones of hyperglycemic mice (Supplementary Fig 4) that were fed a moderately HFD for 16 weeks as measured by qPCR while protein levels did not change compared to ND (Figure 3D-F).

**Figure 3:**
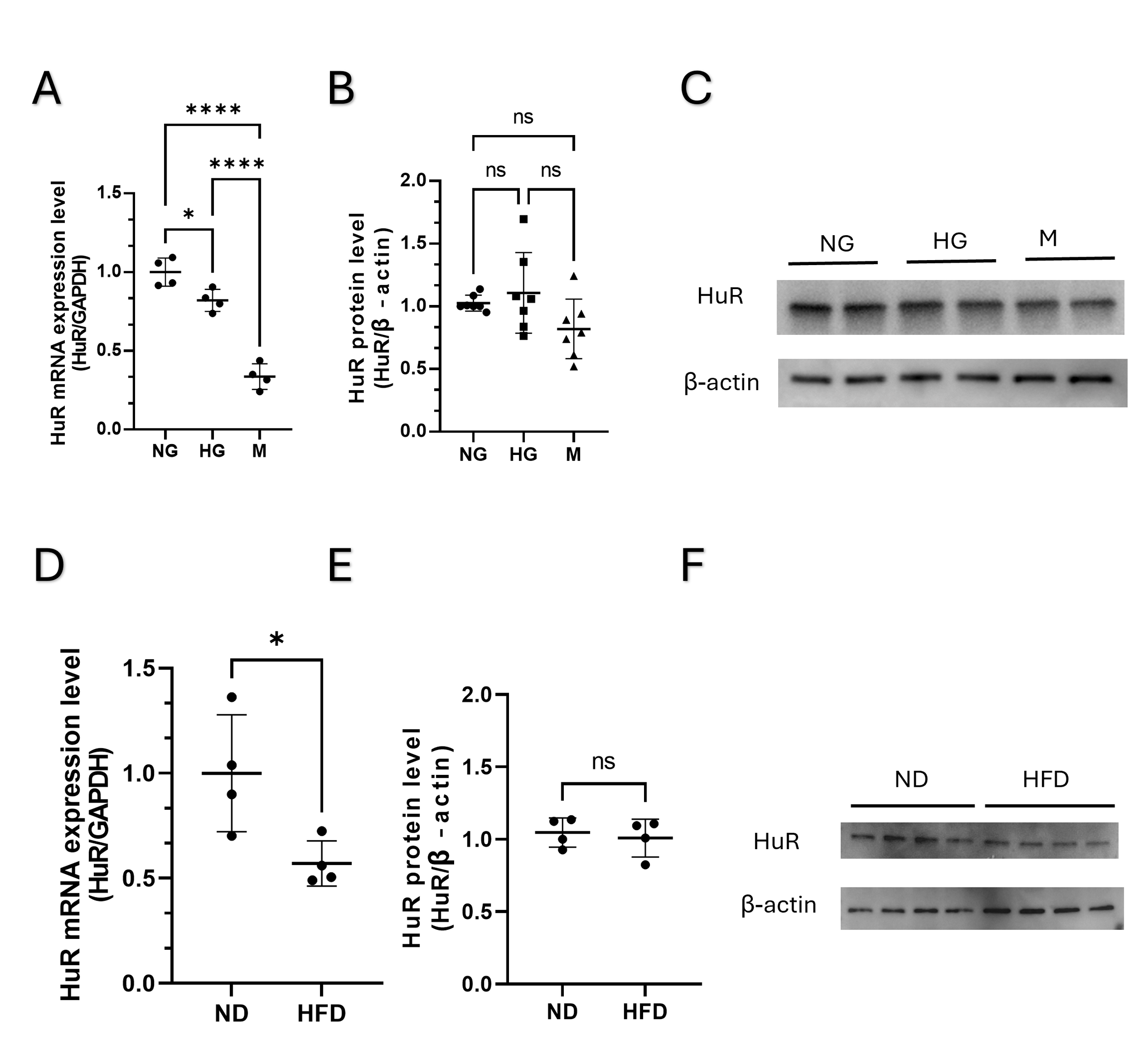
HuR mRNA is reduced in MLO-Y4 cells cultured in HG and in the long bones of HFD fed mice, while maintaining HuR protein levels. **(A)** HuR mRNA level in MLO-Y4 cells under normal glucose (NG), high glucose (HG) and osmotic control mannitol (M). Data are presented as mean ± SD, n=4. *p<0.05, ****p<0.0001. **(B)** HuR protein level in MLO-Y4 cells in normal glucose (NG), high glucose (HG) and osmotic control mannitol (M). Data are presented as mean ± SD, n=7. **(C)** HuR mRNA level in the long bones of high fat diet (HFD) and normal diet (ND) fed mice. Data are presented as mean ± SD, n=4. *p<0.05. **(D)** HuR protein level in the long bones of high fat (HF) and normal diet (ND) fed mice. Data are presented as mean ± SD, n=4. *p<0.05.

### HuR KD reveals transcriptomic and phenotypic changes in MLO-Y4 cells

We knocked down HuR in MLO-Y4 cells (Y^KD^) using two independent shRNA constructs, both of which effectively reduced HuR protein levels (Figure 4A). Despite comparable knockdown efficiency, we observed differences between the two constructs. Construct A (YA^KD^) induced severe cytotoxicity, including a significant loss of cell viability (Figure 4B) and dramatic morphological changes characterized by a severely reduced cytoplasmic volume and near complete loss of dendrites (Figure 4C, D). Additionally, GSEA results from YA^KD^ showed enrichment in DNA damage and cell cycle checkpoint pathways in addition to RNA metabolism and splicing term enrichment espite minimal evidence of differential splicing (Supplementary Figure A, 5B). This pattern is consistent with global stress response rather than specific HuR-dependent regulation. These phenotypic changes suggest potential off-target effects. Construct B (YB^KD^) experienced reduced cell viability and morphological changes though to a lesser extent than YA^KD^, and preserved cell dendrites (Figure 4C, D). To ensure that our results were not confounded by potential off-target effects in construct A, we used YB^KD^ for further analysis, hereafter, the terms HuR KD and YB^KD^ are used interchangeably. RNA-seq analysis of YB^KD^ revealed 129 differentially expressed genes at |Log2FC|≥ 1 and adjusted p value < 0.05 compared to Mock^KD^ including HuR downregulation (Figure 4E). The top 10 enriched pathways in pre-ranked gene set enrichment analysis showed downregulation of extracellular matrix organization and remodeling, including several collagens, matrix metalloproteinases and cathepsins-growth and development pathways, fatty acid and cholesterol biosynthesis pathways, in addition to cell signaling pathways involved in inflammation and cellular homeostasis. Upregulated pathways showed a clear trend towards enrichment in pathways related to quality control of mRNA processing, translation and ribosomal terms. In addition to pathways related to immunity and pathways related to cytoskeleton organization (Figure 4F and Supplementary Table 3). Overall, the transcriptomic profile of HuR KD points to a complex role for HuR in controlling several aspects of cellular homeostasis including cytoskeletal organization, extracellular matrix proteins and a role in regulating mRNA processing and translation.

**Figure 4:**
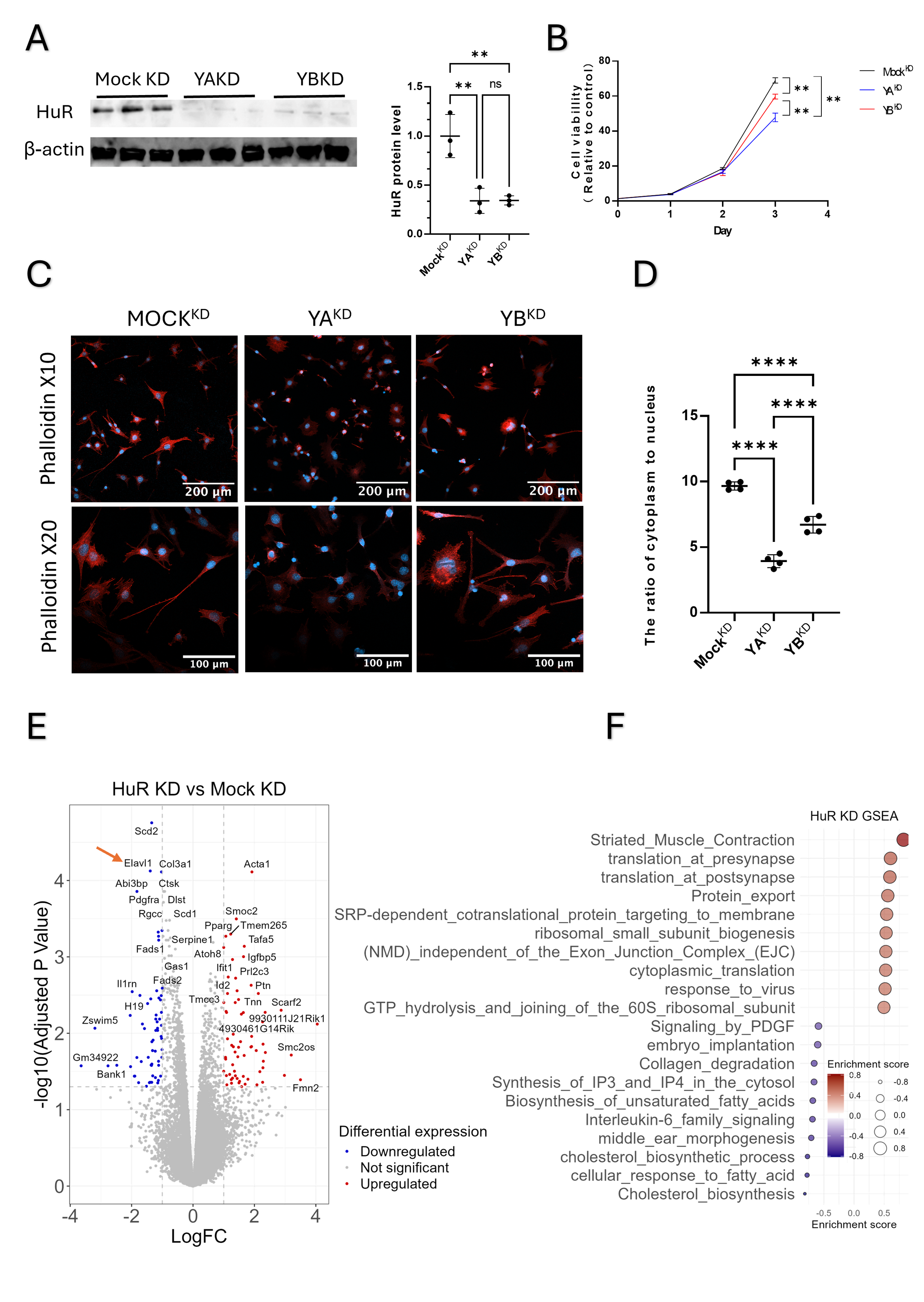
HuR KD confirms functionality in in MLO-Y4 cells. **(A)** Western blot analysis of HuR protein levels after shRNA-mediated knockdown in MLO-Y4 cells. Data are presented as mean ± SD, n=3. **p<0.01. **(B)** CCK cell viability assay. Data are presented as mean ± SD, n=4. **p<0.01. **(C)** Confocal microscopic images of phalloidin staining, Scale bars: 200 µm for the 10× image and 100 µm for the 20× image. **(D)** Cytoplasm to nucleus ratio. The mean of 5 analysed regions (corners and center) of a 35 mm glass bottom dish were used. Data are presented as mean ± SD, n=4. ****p<0.0001. **(E)** Volcano plot comparing the log2 FC and -log10 adjusted P value of RNA-seq data (DEGs) after YB^KD^ in MLO-Y4 cells, red arrow =HuR. n=3. **(F)** Gene set enrichment analysis using eVITTA toolbox after YB^KD^ of DEGs at LogFC 1 and -log10 adjusted p<0.05.

### HuR regulates translational and metabolic stress adaptation splicing programs

We further compared alternative mRNA splicing patterns between YB^KD^ and Mock^KD^. We summarized the output of the AS analysis from junction and exon body counts (JCEC), the total number of JCEC events was 24 thousand events, with 3% showing significance between YB^KD^ and Mock^KD^. Among the AS patterns detected, SE represented the most frequently regulated events, while RI events were the least regulated (Figure 5A). This observation aligns with the earlier HuR motif analysis in primary osteocytes (Supplementary Fig 2 and 3), where RI events showed the least enrichment in HuR motifs around differentially spliced introns. DSGs (Figure 5B, Supplementary Table 4) were used to perform gene ontology (GO) and over representation functional annotation (ORA) following HuR KD. The top 10 scoring cellular components included ribonucleoprotein complexes (RNPs), ribosomes and mitochondria (Figure 5C). ORA analysis results revealed two major clusters of enriched pathways. First, translation-related terms, including rRNA processing, RNA splicing, mRNA surveillance (nonsense-mediated decay), ribosomal subunits, and translational regulation consistent with HuR’s role as an RBP. Second, metabolic or stress-response terms, including autophagy, HIF-1α signaling, and insulin signaling pathways (Figure 5D, Supplementary Table 4).

**Figure 5:**
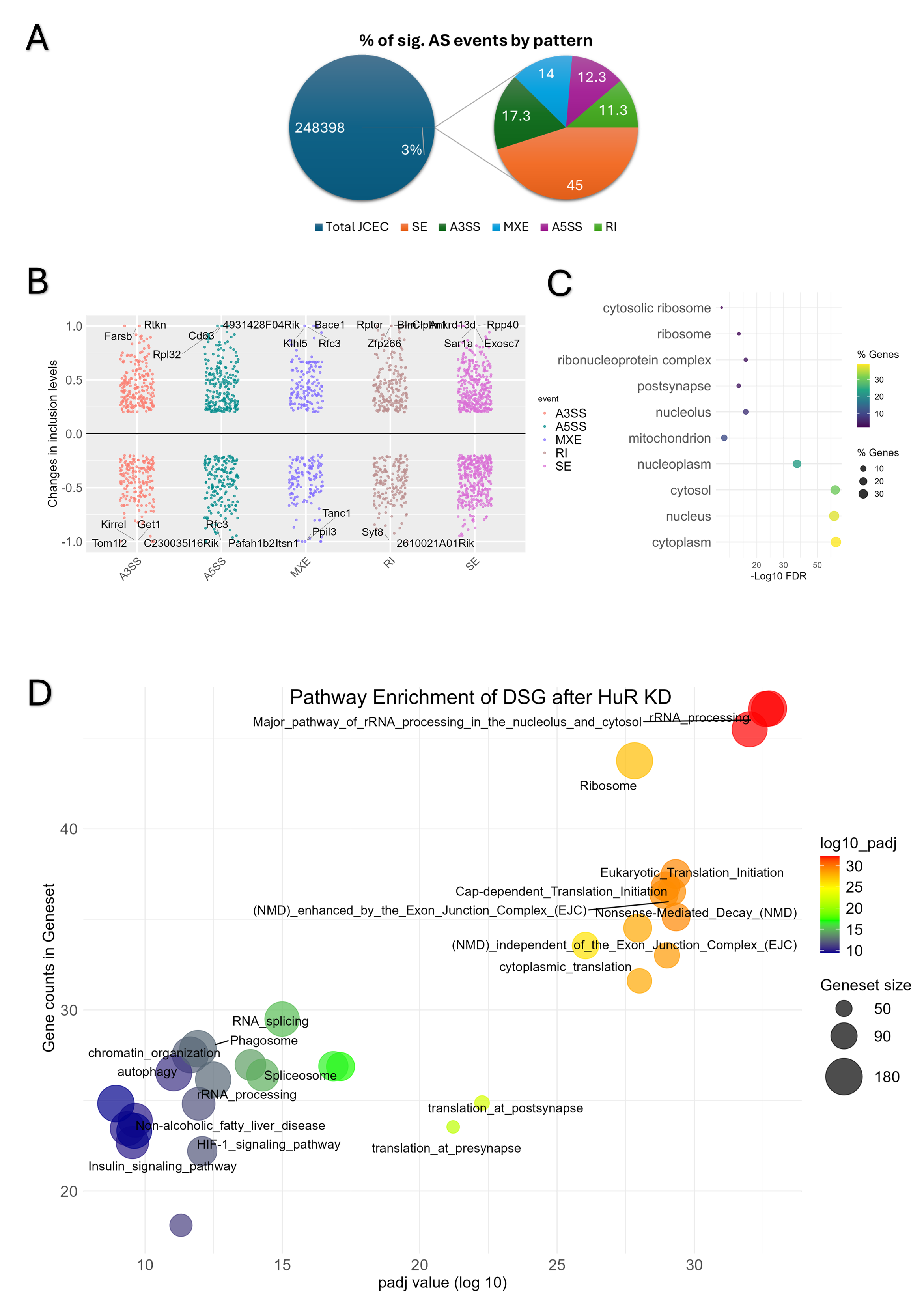
HuR KD induces AS changes in MLO-Y4 cells. **(A)** The parent pie chart shows significant AS events (3%) of the total events at target and junction reads (JCEC) and secondary pie chart showing the percentage of significant AS events of each type (SE, A5SS, A3SS, MXE, RI) from total significant events. **(B)** Scatter plot showing DSGs around AS events in 5 patterns FDR < 0.05 and 0.2 < Δ1 < -0.2. **(C)** Gene Ontology Cellular Component (GOCC) enrichment analysis of DSGs performed using the DAVID Functional Annotation Tool. **(D)** ORA of DSGs after YB^KD^. Also, see Supplementary Table 8.

### HuR KD sensitizes cells to hyperglycemic and oxidative stress induced cell death

Our initial analysis pointed to HuR motif as a top enriched motif around spliced transcripts in HG conditions, therefore we assed the effect of HG on YB^KD^ cells. We cultured cells with NG or HG and M as osmotic pressure control, we used H_2_O_2_ as a positive control. HG induced cell death only in YB^KD^ cells but not in Mock^KD^ after 48 and 72 hours indicating that HuR KD sensitizes cells to hyperglycemic injury. Hydrogen peroxide induced oxidative cell death in both cell lines; but with markedly greater lethality in YB^KD^ compared to Mock^KD^ (Figure 6A). HG is known to induce oxidative stress, therefore, we next examined whether these changes affected redox homeostasis and cellular susceptibility to oxidative stress. We performed DCFDA assay which measures ROS levels, the results indicated a decrease in ROS production under baseline culture conditions in YB^KD^, however, upon treating cells with TBHP, a ROS generating compound, ROS accumulation doubled in YB^KD^ compared to Mock^KD^ (Figure 6B).

**Figure 6:**
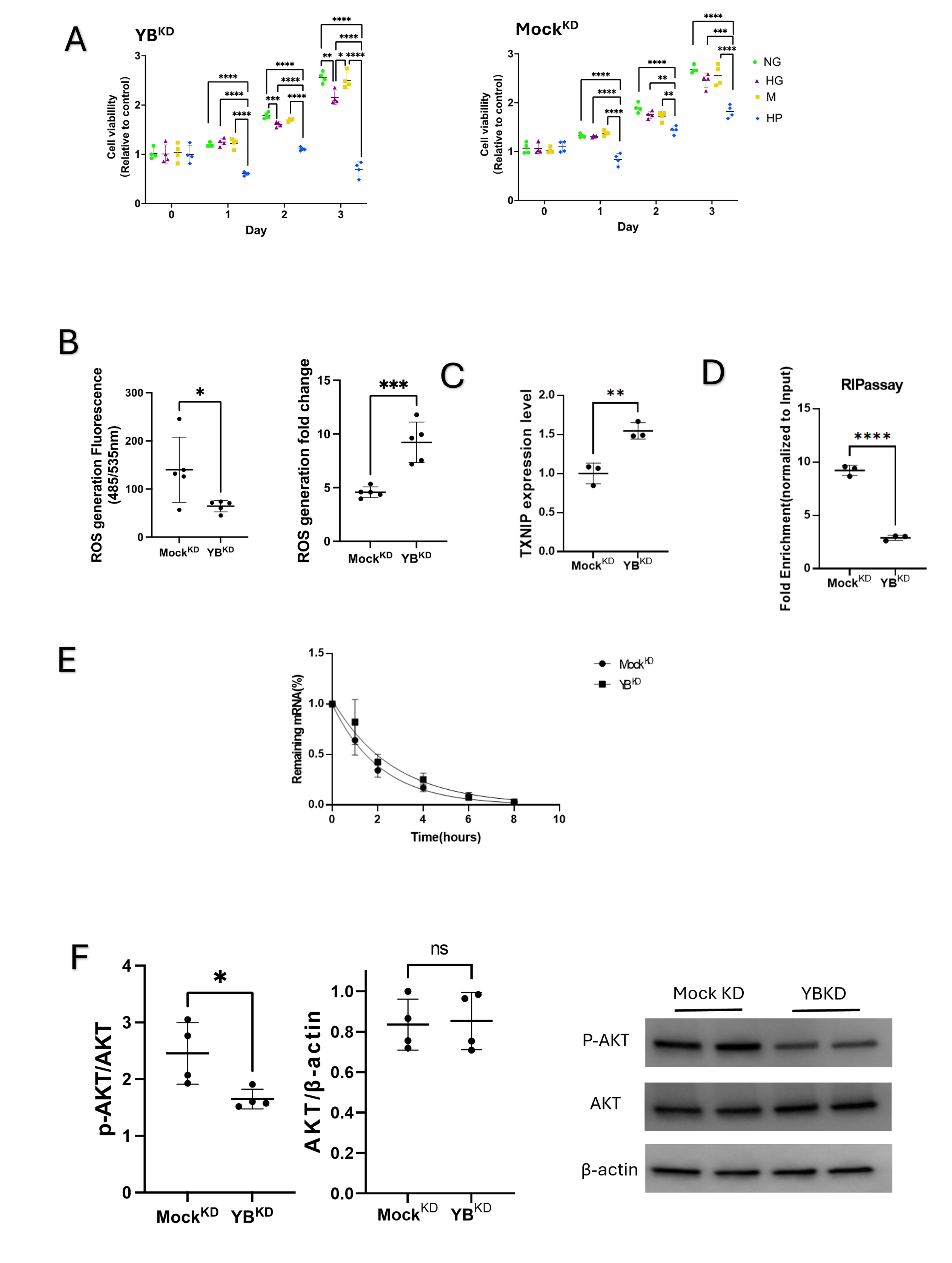
HuR knockdown heightens oxidative stress through enhancing TXNIP stability. **(A)** Cell viability assay in YB^KD^ and Mock^KD^ cells in 5.5mM glucose (NG), 25 mM glucose (HG), mannitol (M) and H_2_O_2_ stimulated cells. Data are presented as mean ± SD, n=4. *p<0.05, **p<0.01,***p<0.001, ****p<0.0001. **(B)** Reactive oxygen species (ROS) measurement. Fold change is calculated after treating cells with TBHP to induce ROS formation. Fold change represents TBHP stimulation/ basal line culture conditions. Data are presented as mean ± SD (n=5 per group; *p<0.05, ***p<0.001) **(C)** TXNIP mRNA level in MLO-Y4 cells after HuR KD. Data are presented as mean ± SD, n=3. **p<0.01. **(D)** HuR RIP-qPCR in MLO-Y4 cells demonstrating TXNIP mRNA enrichment in HuR immunoprecipitates relative to IgG control. Data are presented as relative quantity normalized to input prepresent as mean ± SD, n = 3, ****p < .00001. **(E)** TXNIP mRNA stability assay following actinomycin D treatment. Decay curves were fitted using a one-phase exponential model; half-life increased from 1.47 h in Mock^KD^ to 1.85 h in YB^KD^. Data points represent mean ± SEM (n = 6), p < 0.03 for decay rate constant (K) comparison. **(F)** AKT/ actin and p-AKT/AKT protein levels in YB^KD^ and Mock^KD^ cells. Data are presented as mean ± SD, n=4. *p<0.05.

### HuR depletion disrupts oxidative stress adaptation by stabilizing TXNIP mRNA

To further explore the mechanism by which HuR regulates oxidative stress response, we measured the expression levels of TXNIP. TXNIP mRNA levels increased after HuR knockdown (Figure 6C). We then performed RNA immunoprecipitation (RIP), which showed that HuR-specific antibody effectively co-precipitated TXNIP mRNA compared to negative IgG control and precipitated 70% more TXNIP mRNA than Mock^KD^ confirming direct interaction (Figure 6D). We then quantified TXNIP mRNA decay after actinomycin D treatment; HuR KD increased TXNIP stability (Δt½ = 1.470 → 1.846 hours), an almost 25% increase in mRNA half-life. Together, HuR binds TXNIP mRNA and limits its stability, providing a mechanistic link between HuR loss and heightened oxidative stress sensitivity.

### HuR regulates insulin sensitivity

Our DSG analysis pointed to insulin signaling term enrichment, and insulin is a known suppressor of TXNIP expression.^34^ Therefore, we investigated the functionality of our DSG analysis (Figure 5D). We assessed whether HuR KD altered insulin responsiveness. HuR KD exhibited a reduced response to insulin stimulation, as evidenced by decreased AKT phosphorylation (p-AKT) in Figure 6F.

### HuR KD induces mitochondrial dysfunction

Our DSG analysis revealed enrichment of mitochondrial-related terms, and HuR KD cells showed both heightened susceptibility to ROS-induced death and increased ROS accumulation. Because mitochondrial respiration is a primary source of cellular ROS, these findings prompted us to examine mitochondrial mass and function. To quantify mitochondrial density, we used Mitotracker CMXRos Red, a selective mitochondrial dye. Mitochondria localized around the nuclei and extended to dendrites, and we observed reduced mean fluorescence intensity in YB^KD^ cells compared to Mock^KD^ (Figure 7A, B). Next, we measured mitochondrial function using Seahorse XF Cell Mito-Stress test (Figure 7C). YB^KD^ cells exhibited reduced mitochondrial respiration. Basal OCR, maximal respiration, spare respiratory capacity, coupling efficiency and ATP production were reduced. Proton leak remained unchanged, suggesting that the mitochondrial membrane is intact. The observed reduction in OCR also extended to non-mitochondrial respiration, suggesting that the decline in oxygen consumption is partly attributable to reduced activity of non-mitochondrial oxygen-consuming enzymatic reactions (Figure 7D).

**Figure 7:**
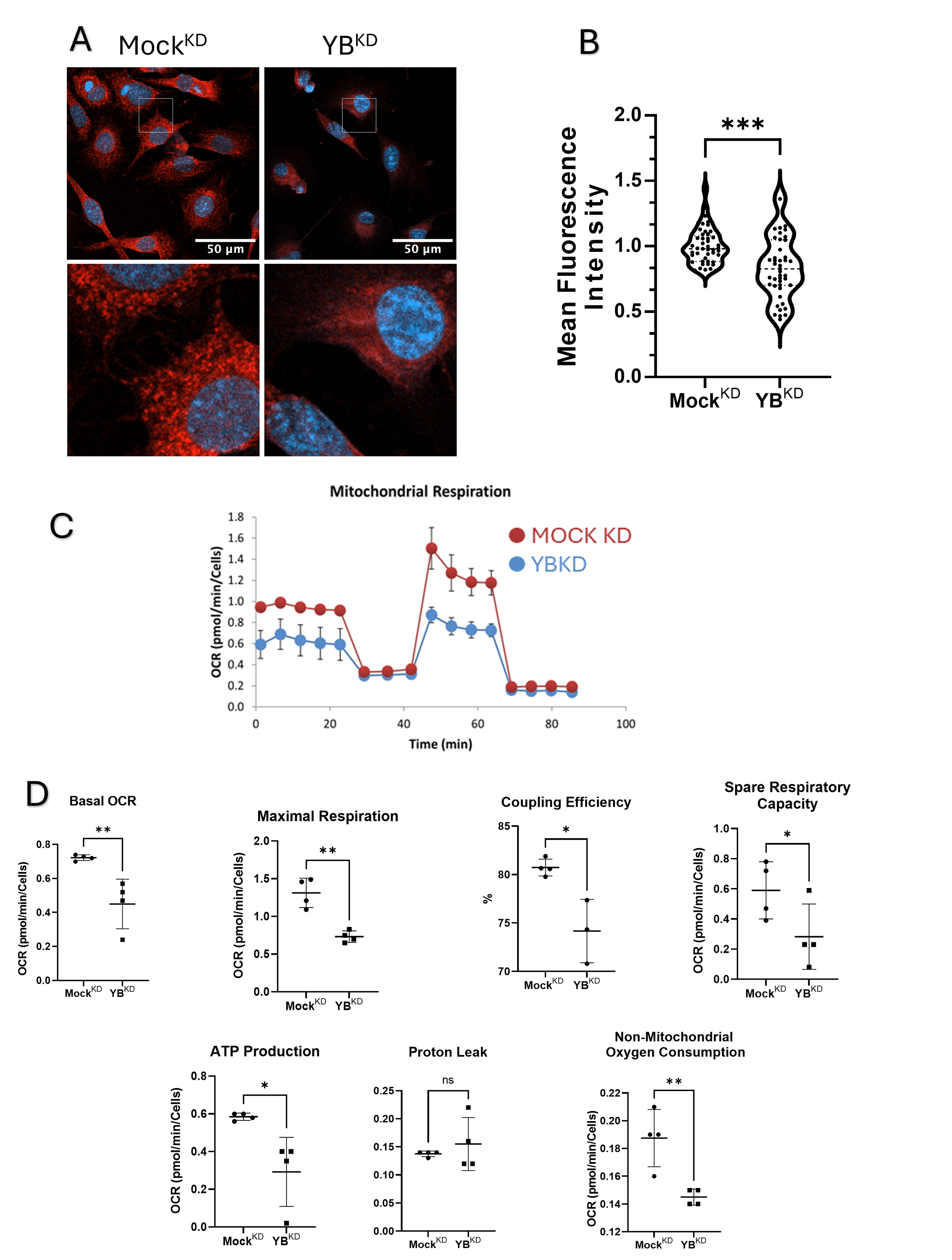
HuR knockdown induces mitochondrial dysfunction. **(A)** Fluorescent microscopic images of Mitotracker CMXRos Red staining mitochondria. **(B)** Mean fluorescent intensity (MFI) of Mitotracker CMXRos Red staining mitochondria. Data are presented as mean ± SD, n=4. ***p<0.001. **(C)** Seahorse XF Cell Mito-Stress showing oxygen consumption rate (OCR) in pmol/min. **(D)** Parameters of mitochondrial respiration. Data are presented as mean ± SD, n=4. *p<0.05, **p<0.01)

### HuR KD disrupts translational signaling while preserving global protein output

Our DSGs analysis revealed significant enrichment in pathways related to translational control, therefore, we investigated several key translational signals. We assessed the phosphorylated status of the mechanistic target of rapamycin (mTOR), a central mediator of mRNA translation and protein synthesis. mTOR phosphorylates two key immediate targets, the p70S6 kinase (p70S6K) and eukaryotic initiation factor 4E-binding protein 1 (4E-BP1).^35^ Total levels of p70S6K and 4E-BP1 were reduced in HuR KD cells. However, their phosphorylated forms, p-p70S6K and p-4E-BP1, increased, resulting in increased phosphorylated-to-total protein ratios (Figure 8A, B). We next examined mitogen activated protein kinase-interacting serine/threonine kinase 1 (MAPK-MNK1) axis, which predominantly regulates cap dependent translation initiation, the predominant form of translation in eukaryotic cells. Phosphorylated MNK1 decreased, whereas total MNK1 levels remained unchanged (Figure 8C). Consistently, upstream of MNK1, the extracellular signal-regulated kinases1/2 (ERK1/2) showed reduced levels of both phosphorylated and total protein (Figure 8D). Furthermore, these pathways converge on the eukaryotic initiation factor 4E (eIF4E), a direct substrate of MNK1 and 4E-BP1. In HuR KD cells, both total and phosphorylated levels were reduced, leading to a lower p-eIF4E/eIF4E ratio (Figure 8E). To determine whether these molecular changes affected overall protein translation, we performed a puromycin incorporation assay to measure nascent protein synthesis rate. Despite the pronounced alterations in these translational regulators, no significant differences in global protein synthesis rates were detected suggesting maintenance of translational output through the compensatory activation of mTOR1 (Figure 8F).

**Figure 8:**
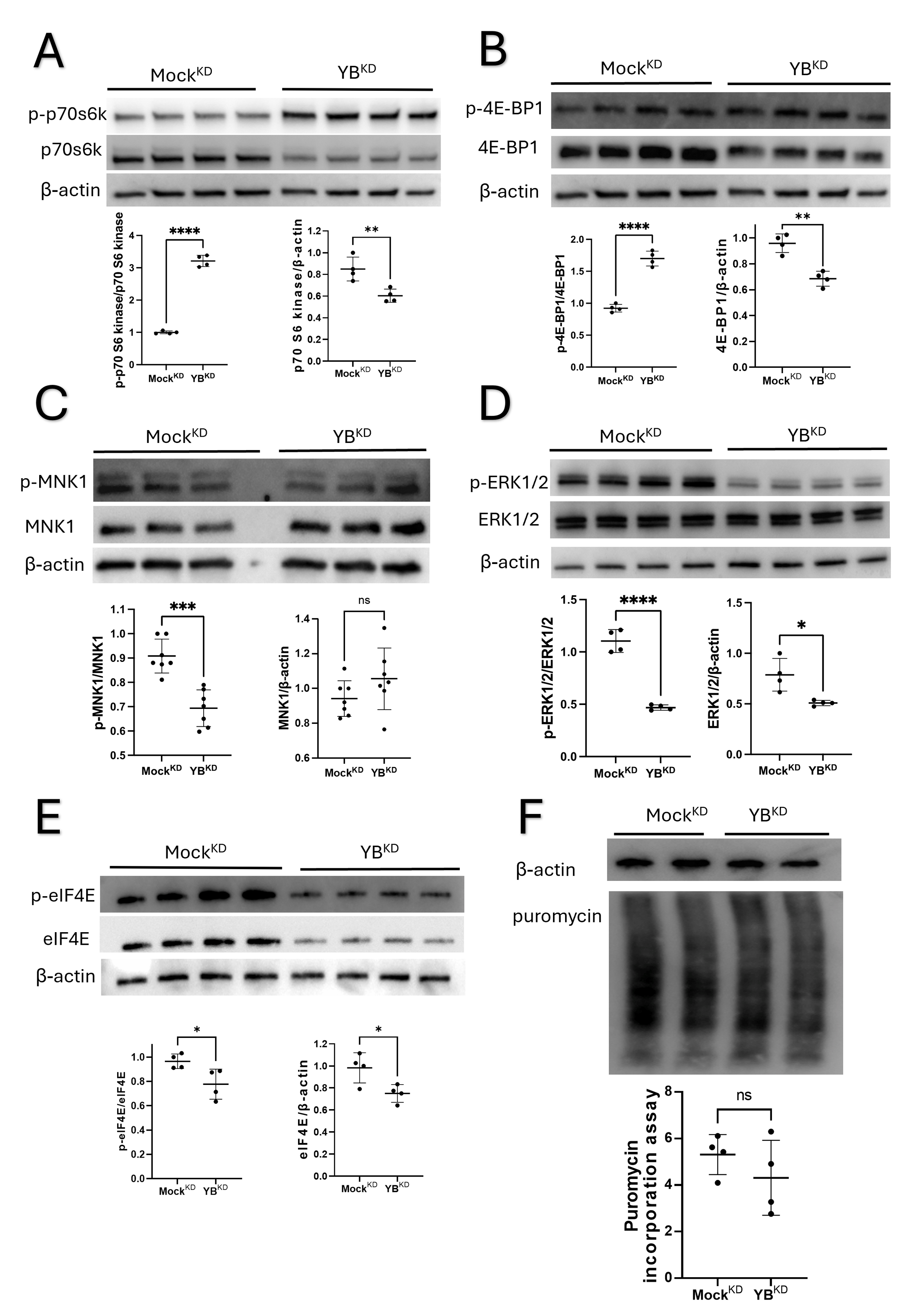
HuR knockdown induces translation dysregulation and preserves global protein synthesis rate. **(A)** Western blot of p-p70S6K and p70S6K protein levels. n=4. **(B)** Western blot of p-4E-BP1 and 4E-BP1 protein levels. n=4. **(C)** Western blot of p-MNK1 and MNK1 protein levels. n=7. **(D)** Western blot of p-ERK1/2 and ERK1/2 protein levels. n=4. **(E)** Western blot of p-eIF4E and eIF4E protein levels. n=4. **(F)** Western blot of puromycin incorporation assay. n=4. Data are presented as mean ± SD of 2 or more independent blots (*p<0.05, **p<0.01, ****p<0.0001)

## Discussion

RNA fate and function are largely determined by its interaction with RBPs which recognize consensus RNA binding motifs or structural domains to modulate RNA splicing, stability, localization and translation, known as post-transcriptional gene regulation. Post-transcriptional gene regulation ultimately shapes physiological and adaptive cellular response to stress. HuR, is a ubiquitously expressed RBP that contains three RRM domains that mediate binding to adenine- and uridine-rich elements (AREs) predominantly located within the 31 untranslated regions (31UTRs) of target transcripts. Through these interactions, HuR is best known for modulating its target mRNA stability and modulating their alternative splicing ^36,37^.

HuR has emerged as a key modulator of metabolic disease, and its role appears to vary across cells, tissues and disease models. In adipose tissue, HuR levels are decreased in diabetic and HFD-fed mice, and adipose-specific HuR knockout predisposes mice to HFD-induced obesity, insulin resistance, and glucose intolerance.^22^ In the liver, HuR expression is similarly reduced in obesity-associated steatosis, yet its knockout aggravated hepatic steatosis in some models,^38^ and in others worsened insulin resistance and glucose intolerance.^23^ Conversely, increased HuR expression has been observed in diabetic human heart and rat retina, suggesting context-dependent upregulation.^39,40^

These context-dependent discrepancies also manifest in the skeletal system. In ovariectomy-induced osteoporosis, HuR levels decreased, and its overexpression mitigated bone loss, while silencing impaired osteoblastic differentiation in MC3T3-E1 cells.^25^ In contrast, a diabetic mouse model exhibited elevated HuR mRNA levels, and knockdown improved glucose homeostasis and bone structure.^26^ Similarly, high glucose exposure increased HuR expression in MC3T3-E1 cells, where silencing attenuated apoptosis and enhanced osteogenic differentiation.^26^ Collectively, these reports underscore that HuR responds to metabolic perturbations in a tissue- and condition-specific manner, possibly due to the effect being enacted by the target RNA affected rather than HuR abundance itself.

Our transcriptome wide analysis in osteocytes exposed to high glucose, revealed minimal differential gene expression, whereas alternative splicing changes were prominent, and regulated a wide array of cellular programs including ECM proteins, translation, and cytoskeletal organization. These findings prompted us to investigate post-transcriptional regulation under high glucose. To probe the regulatory factors underlying these splicing alterations, we used the differentially spliced gene sets to identify RBP motif enrichment which identified several RBPs including HuR, ZC3H14, and HNRNPL as recurrently enriched around differentially spliced exons. HuR emerged as a prominent candidate, appearing in multiple positions across various splicing events. Therefore, we examined HuR expression at both the mRNA and protein levels in osteocytic MLO-Y4 cells cultured in HG and in the bones of HFD mice. Despite a significant reduction in HuR mRNA, protein levels remained unchanged, suggesting post-transcriptional buffering. Interestingly, mannitol-treated cells used to control osmotic stress exhibited a similar pattern, with decreased HuR mRNA but stable protein levels, further implicating regulatory mechanisms that maintain protein abundance under stress. Previous studies have shown that HuR protein levels remained stable under hypotonic stress, while its subcellular localization shifted from the nucleus to the cytoplasm.^41^ Although we did not directly assess HuR localization in our study, it is conceivable that similar stress-induced relocalization occurred in our cells, potentially contributing to the observed effect on mRNA and protein levels. Our observation is also consistent with evidence that mRNA abundance does not reliably predict protein levels, due to translational control and buffering mechanisms.^42^ Given HuR’s central role in maintaining cellular homeostasis, its expression may be preserved through enhanced translational efficiency or altered localization to sustain functional output. However, this hypothesis remains speculative and warrants further investigation using approaches such as ribosome profiling or spatial proteomics.

The dynamics of HuR expression under hyperglycemic stress prompted us to investigate its role in osteocytes. HuR KD in MLO-Y4 cells exhibited profound differential gene expression, including reduced expression of genes involved in extracellular matrix remodeling consistent with its reported role in maintaining several ECM proteins including collagens and matrix metalloproteinases.^43^ DEGs also showed downregulation in lipid metabolism and cytoskeletal organization reflecting cellular morphological changes. Consistent with that phalloidin staining showed rounding and loss of dendrites after HuR KD consistent with GSEA. Enriched pathways were clearly biased towards mRNA quality control and mRNA translation.

DSGs in HuR KD cells showed enrichment in ribosomal and mitochondrial terms as well as terms related to the translational machinery, such as rRNA processing, RNA splicing, and translation-dependent mRNA surveillance pathways, including NMD, indicating an increased aberrant mRNAs and mRNA degradation.^44^ These findings highlight that HuR KD induces substantial splicing alterations that correlate well with transcriptional alterations. Furthermore, AS analysis revealed significant enrichment in stress-related pathways including autophagy and HIF-1α signaling as well as key metabolic pathways such as insulin signaling and non-alcoholic fatty liver disease. Notably, these pathways were not detected through differential gene expression analysis, highlighting AS as a critical post-transcriptional mechanism of gene regulation. These findings emphasize the added resolution provided by splicing analyses, enabling the identification of regulatory changes that are not apparent at gene abundance levels alone.

HuR KD increased the sensitivity of osteocytic cells to glucose-dependent oxidative stress and displayed elevated ROS levels following exposure to oxidative agents supporting a protective role for HuR in maintaining redox balance. Interestingly, under baseline conditions, HuR KD led to a reduction in ROS levels. TXNIP, is a glucose-inducible inhibitor of thioredoxin (TRX), a ROS scavenger that protects against diabetic bone loss.^45^ Our results indicated a direct link between TXNIP mRNA and HuR, as HuR coprecipitated with TXNIP, and upon HuR depletion TXNIP mRNA levels and half-life increased indicating that HuR destabilizes TXNIP mRNA. Our results are also corroborated by previous data reporting TXNIP among 4874 consensus genes that directly bind HuR ^37^.

Concomitant with these results, DSG analysis showed an enrichment in insulin signaling related terms. To validate we probed insulin responsiveness. HuR KD impaired insulin signaling, as evidenced by reduced AKT phosphorylation in response to insulin stimulation. Insulin reportedly suppresses TXNIP expression in insulin responsive cells.^34^ Collectively, HuR loss elevates TXNIP via two coordinated mechanisms, reduced mRNA decay and attenuated insulin-dependent transcriptional downregulation, thereby, increasing oxidative-stress sensitivity under high glucose.

Given that mitochondria are the primary source of cellular ROS, we assessed mitochondrial mass and function. HuR KD reduced mitochondrial content, as estimated by Mitotracker staining, and impaired mitochondrial respiration, evidenced by decreased OCR, maximal respiration, and coupling efficiency. ATP production was also diminished, indicating reduced oxidative phosphorylation (OxPhos), while proton leak remained unchanged, suggesting that mitochondrial membrane integrity was preserved. The decrease in ROS under baseline conditions may be explained by reduced mitochondrial activity and total oxygen consumption by cells reducing ROS generating capacity. However, the heightened vulnerability to exogenous oxidative stress under HG and ROS-inducing conditions highlights a failure of adaptive antioxidant response in HuR KD cells. Collectively, these findings underscore HuR’s role in maintaining cell viability under HG conditions and maintaining redox homeostasis and mitochondrial function.

Given the enrichment of AS events in pathways related to mRNA translation and protein synthesis, we examined whether these changes were functionally reflected in translational signaling. Eukaryotic translation can be initiated via two mechanisms, cap-dependent and cap independent translation. Cap dependent translation is the predominant form and cap dependent translation initiation is the rate limiting step in translation which depends on eIF4E availability.^46^ eIF4E binds to the m^7^G cap located at the 5’ end of most mRNAs where it recruits other initiation factors preparing the mRNA for ribosomal loading and scanning for the translation initiation codon.^46^ The activity of eIF4E is regulated via two mechanisms. First, it is sequestered by the inhibitory 4E-BP1 which releases eIF4E upon its phosphorylation by mTORC1 kinase.^35^ Second, available eIF4E is phosphorylated by MNKs which serves to modulate selective mRNA translation. Notably, phosphorylation of eIF4E by MNKs is not required for global mRNA translation, conversely, mTORC1 inhibition leads to near complete inhibition of mRNA translation.^47,48^ This aligns well with our data where HuR KD preserved global protein output possibly due to enhanced mTORC1 activity and inhibition of 4E-BP1 which serves to release eIF4E for translation initiation.

Our data revealed reduced total levels of p70S6K and 4E-BP1, yet their phosphorylated forms were elevated, resulting in increased phospho-to-total ratios—suggesting relative activation of mTORC1 kinase. In contrast, the MNK1-eIF4E axis was suppressed: phosphorylated MNK1 and its upstream kinase ERK1/2 were reduced, and both total and phosphorylated eIF4E levels were diminished. Despite these alterations, global protein synthesis rates remained unchanged. The preservation of global protein output despite disrupted signaling in key translational pathways suggests a shift toward biased mRNA translation, where the cell prioritizes the synthesis of specific mRNAs. Such reprogramming may represent an adaptive response to preserve proteostasis under energetic and metabolic stress. Although we did not measure selective mRNA translation it is plausible that a shift towards increased translation of mRNAs related to cell survival has taken place in HuR KD cells.

In conclusion, our study elucidates the role of HuR in osteocyte biology, particularly in regulating AS, translation, mitochondrial function and redox homeostasis. HuR seems to play a protective role against high glucose and oxidative cell death. Our findings reveal that HuR is indispensable for maintaining cellular and metabolic homeostasis in osteocytes, particularly under HG conditions. Our data provides a mechanistic understanding of HuR’s role in cellular homeostasis and has broader implications in metabolic bone diseases.

## Materials and Methods

### 1. Primary cells and cell lines

Primary osteocytes were isolated from neonatal calvariae of C57BL/6-Tg (Dmp1-Topaz)1Ikal/J using serial enzymatic digestion with collagenase and EDTA followed by fluorescent activated cell sorting (FACS) as described previously.^49^ After isolation, cells were cultured at standard conditions (37 LJ1, 5% CO_2_) overnight in alpha-minimum essential medium (1-MEM; Wako Pure Chemical Industries, Osaka, Japan) containing 10% fetal bovine serum (FBS; Biowest, Nuaillé, France), 100IU/mL penicillin G, 100 μg/mL streptomycin (1% P/S) containing 5.5 mM glucose (standard medium). After that, the media was replaced with fresh 5.5mM glucose (normal glucose, NG), 25 mM glucose (Sigma-Aldrich, St. Louis, MO, USA) (high glucose, HG) containing media or 5.5 mM glucose supplemented with 19.5 mM mannitol (osmotic control, M) (Sigma-Aldrich) for 3 days. MLO-Y4 cells (AddexBio Technologies) were maintained at 37 LJ1, 5% CO_2_ in standard medium and kept at 70-80% confluency and passaged as needed. Standard medium was changed to NG, HG or M media for subsequent experiments.

### 2. Mice and feeding experiments

12-week-old male C57BL/6J mice were purchased from CLEA Japan (Tokyo, Japan). Mice were acclimated for 4 weeks in specific pathogen-free (SPF) conditions under a 12-h light/dark cycle with ad libitum feeding (Research Diet, D12450H, Japan) and at 16 weeks the mice were randomly allocated to a 10% Kcal from fat termed normal diet (ND) (Research Diet, D12450H, Japan) or 45% Kcal from fat termed high fat diet (HFD) (Research Diet, D12451, Japan) for 16 weeks to induce chronic hyperglycemia. The mice were tested for fasting blood glucose and glucose tolerance at 4 and 8 weeks of diet and one day before collecting the bones at 16 weeks of diet with 5 mice per group. After sacrifice, the long bones (femur, tibia, humerus) were excised. The muscles and soft tissues were carefully removed, and the bones were scraped using a scalpel. The epiphyses were cut and the bone marrow flushed in phosphate-buffered saline (PBS) x1 at 9000 xg, 4 LJ1 three times. The bones were immediately frozen in liquid nitrogen and stored at -80 LJ1 for RNA and protein extraction. All procedures involving animals were performed in accordance with the ARRIVE guidelines and were approved by the Regulations for Animal Experiments and Related Activities at Tohoku University (2021DnLMO-007-04).

### 3. Blood glucose measurement and glucose tolerance test

Fasting blood glucoseand glucose tolerance were assessed after a 6-hour fast (8:00 AM–2:00 PM) using a rodent specific glucometer with compatible LG sensors (LabGluco Funakoshi., Tokyo, Japan). Approximately, 5 μL or less of blood was drawn from a tail tip clip at 0, 15, 30, 60, and 90 minutes.

### 4. Short hairpin RNA knockdown

Two short hairpin RNA (shRNA) sequences targeting mouse HuR/ELAVL1 mRNA were designed using VectorBuilder’s shRNA Target Design tool. An empty backbone sequence was used as a non-targeting control. The shRNA oligonucleotides were ordered and synthesized by FASMAC Co., Ltd. (Kanagawa, Japan). Oligos were annealed and ligated and cloned into pLKO.1 puro vector (Addgene, Plasmid# 8453). Plasmids were then extracted using Nucleospin plasmid DNA purification kit (Macherey-Nagel, Düren, Germany) followed by validating the insertions using colony-PCR confirming sequences by Sanger Sequencing (Eurofins Genomics, Tokyo, Japan). HEK293T cells were co-transfected with shRNA plasmids, lentiviral packaging plasmid psPAX2 (Addgene, Plasmid#12260) and VSV-G envelope expressing plasmid pMD2.G (Addgene, Plasmid#12259) using Lipofectamine 3000 (Thermo Fisher Scientific, Waltham, MA, USA). Viral particles were collected after 48 and 72 hours, centrifuged at 1500 xg for 5 minutes and supernatant was filtered through a 0.45 µm PVDF filter. Viral particles were stocked at 4 LJ1 for immediate use or -80 LJ1 for long term storage. MLO-Y4 cells were grown to 80% confluency in standard medium, then media was changed to transduction medium containing standard medium and lentiviral particles containing supernatant at 4:1 (by volume) and 8 µg/mL polybrene (Sigma-Aldrich, St. Louis, MO, USA) for 24 hours. Transduction medium was then replaced with standard medium and incubated for 6 hours then puromycin was added at (2 µg/mL) and transfected cells were selected for 7 days before passaging. Construct sequences of control Mock knockdown (Mock^KD^), construct A knockdown (YA^KD^) and construct B knockdown (YB^KD^) in Supplementary Table 5. A and B denote 2 separate constructs and (Y) denotes that MLO-Y4 cells were used for knockdown.

### 5. RNA sequencing

Total RNA was extracted using Qiagen RNeasy kit (QIAGEN, Hilden, Germany) with a DNase digestion step (QIAGEN) as per manufacturer’s instructions. The samples were processed by Azenta, Japan as follows: samples were screened and quantified on an Agilent 4200 Tapestation with RNA ScreenTape. The resultant RIN (RNA integrity number) was 9.9 or 10 for all samples. mRNA was enriched using NEBNext Poly(A) mRNA Magnetic Isolation Module (NEB) and RNA library preparation was done NEBNext Ultra II Directional RNA Library Prep Kit (Illumina E7760). Samples were pooled on an Illumina Novaseq to obtain paired end 150bp reads with a target depth of 50 million reads per sample. FASTQ files were then used for bioinformatics analysis. All sequencing experiments were done in triplicates.

### 6. RNA sequencing analysis

Quality control of raw FASTQ files was conducted using FastQC. Adapters and low-quality reads were removed using Trimmomatic.^50^ Trimmed reads were aligned to the mouse reference genome (M35, GRCm39) using the splice-aware aligner HISAT2.^51^ The resulting BAM files of mapped reads were quantified against gene features using FeatureCounts with default parameters.^52^ Differentially expressed genes (DEGs) were identified using Limma-Voom,^53^ applying TMM normalization and defining DEGs as those with |log₂ fold change| ≥ 1 and adjusted p value < 0.05.

### 7. AS and RBP motif analysis

Aligned BAM files were used to conduct AS analysis using the rMATs-turbo suite.^32^ Statistical significance of AS events was considered at FDR < 0.05 and |Ψ| > 0.2 to filter in differentially spliced genes (DSGs). Splice motif analysis was conducted using rMAPs2.^33^

### 8. Gene ontology and functional enrichment analysis

DSGs were used for functional enrichment analyses. The output of rMATs was piped into rMAPs to generate RBP motif maps.^33^ DEGs and DSGs were analyzed for gene ontology (GO) and functional enrichment analysis using either David GO functional annotation tools or eVITTA’s pre-ranked gene set enrichment analysis (GSEA) or overrepresentation analysis (ORA) toolbox.^54^ Figures in this publication were made using bespoke scripts in Rstudio 4.4.0 ggplot or ggplot2 packages and extensions.

### 9. Cell viability assay

In a 96-well plate, 1×10^4^ Mock^KD^, YA^KD^ or YB^KD^ were seeded in each well. Cells were cultured in each well with 200 μL of α-MEM supplemented with 10% FBS and 1% P/S. Cell viability was measured at 0, 24, 48, and 72 hours. The plate was then incubated for 2 hours at 37°C with each well containing 100 µL of α-MEM and 10 µL of Cell Counting Kit-8 solution (Dojindo Laboratories, Kumamoto, Japan). Absorbance was measured at 450 nm using a microplate reader (Sunrise™ Remote, Tecan, Männedorf, Switzerland).

### 10. Real-time polymerase chain reaction (qPCR)

RNeasy kit (QIAGEN) was used to extract total RNA from cells according to manufacturer’s instructions and Trizol-chloroform was used to extract RNA from bones. 1 femur was immersed in 1 mL Trizol and pulverized 5 times at 5000 RPM speed setting for 60 seconds each in TOMY Micro Smash MS 100 Cell Disruptor. Total RNA was used to synthesize cDNA using the SuperScript® IV First-Strand Synthesis System (Invitrogen, Carlsbad, CA, USA) according to manufacturer’s instructions. mRNA expression values were measured using the CFX96 Touch Real-Time PCR Detection System (Bio-Rad).

### 11. RNA Immunoprecipitation (RIP) assay

RIP assay was performed using the EZ-Magna RIP kit (Millipore, Billerica, MA) according to the manufacturer’s instructions. Briefly, cells at 80% confluency were scraped off and lysed in RIP lysis buffer provided in the kit. Then 100 µL of whole cell lysates were incubated with RIP buffer containing magnetic beads conjugated with anti-HuR antibody (or normal IgG as negative control, Anti-SNRNP70 as positive control). After washing, RNA–protein complexes were eluted and digested with proteinase K, and the RNA was purified using the kit reagents. Purified RNA was subjected to cDNA synthesis and qPCR analysis as described in the RT-qPCR section.

### 12. mRNA stability assay

To estimate mRNA half-life, nascent transcription was blocked with Actinomycin D (ActD, final 101µg/1mL). At t1=10, 1, 2, 4, 6, 8 h after addition, cells were rapidly washed with ice-cold PBS and lysed in TRIzol and RT-qPCR was done as described. mRNA decay rate was measured by non-linear regression curve fitting (one phase decay) using GraphPad Prism using the following parameters: goodness of fit was quantified with R^2^, using ordinary fit at confidence level 95%.

### 13. Insulin response tests

Mock^KD^ or YB^KD^ cells were seeded in α-MEM supplemented with 10% FBS and 1% P/S until reaching 80% confluency. The media were then replaced with α-MEM containing 1% P/S and 1% FBS for 3 hours followed a switch to the same media with or without 100 nM insulin (Sigma-Aldrich). After a 30-minute incubation period, cells were harvested and protein extracted using radioimmunoprecipitation (RIPA) assay buffer (Millipore, Burlington, MA, USA) containing 1% protease and phosphatase inhibitor cocktail (Thermo Fisher Scientific) on ice for 20 minutes; insoluble material was separated by centrifugation at 14000 xg. Total protein was collected for western blot analysis.

### 14. Reactive oxygen species detection assay

Mock^KD^ or YB^KD^ (2.5 × 10^4^ cells per well) were seeded in a black, clear bottom 96-well microplate overnight. Intracellular reactive oxygen species (ROS) levels were measured using the DCFDA/H2DCFDA-Cellular ROS Assay Kit (Abcam, Cambridge, UK) according to manufacturer’s instructions. Cells were stained with 20 μM DCFDA in HBSS for 45 minutes at 37°C. Mock^KD^ or YB^KD^ cells treated with 50µM TBHP for 4 hours were used as positive control. DCF fluorescence intensity was measured using a fluorescence plate reader (Flex Station 3, Molecular Devices, San Jose, CA, USA) with excitation at 485 nm and emission at 535 nm.

### 15. Western blot

Cells were lysed using RIPA assay buffer (Millipore, Burlington) containing 1% protease and phosphatase inhibitor cocktail (Thermo Fisher Scientific) on ice for 20 minutes; insoluble material was separated by centrifugation 14000 xg. Protein from bones was extracted in RIPA buffer and processed similar to RNA extraction above. Total protein was quantified using Pierce™ BCA protein assay kit (Thermo Fisher Scientific). To prepare for SDS-PAGE gel electrophoresis, the protein was treated with β-Mercaptoethanol (Bio-Rad Laboratories) and Laemmli sample buffer (BioRad Laboratories) 3:1 and denatured at 95°C for 5 minutes. Equal protein amounts were loaded onto 4–15% Mini-PROTEAN TGX Precast Gels (Bio-Rad Laboratories) and transferred to a Trans-Blot Turbo Transfer System (Bio-Rad Laboratories) and then blocked with Block-Ace (DS Pharma Biomedical, Osaka, Japan) 1 hour at room temperature (RT). Primary antibodies were incubated overnight at 4°C. The membranes were incubated with horseradish peroxidase-conjugated anti-rabbit antibody (Cell Signaling Technology, Danvers, MA, USA) at a dilution of 1:5,000 or anti-mouse antibody (GE Healthcare, Chicago, IL, USA) at a dilution of 1:10,000 for 1 hour at RT after being washed in tris-buffered saline with Triton X-100 (TBS-T). Bound antibodies were detected with SuperSignalWest Femto Maximum Sensitivity Substrate (Thermo Fisher Scientific) and a FUSION-FX6 EDGE Chemiluminescence Imaging System (Vilber Lourmat, Collégien, France). Primary antibodies are in Supplementary Table 5.

### 16. Confocal microscopy

1 × 10^5^ Mock^KD^, YA^KD^ or YB^KD^ cells were seeded in 35mm glass bottom dish overnight. Cells were fixed with 4% formaldehyde for 15 minutes, then washed 3 times with x1 PBS. Cells were permeabilized using 0.1% Triton X-100 for 10 minutes and blocked with 1.5% bovine serum albumin (BSA) for 30 minutes at RT. Cells were incubated with Alexa Fluor 568 Phalloidin (Invitrogen) for 1 hour at RT. Afterwards, cell nuclei were stained with 4’,6-diamidino-2-phenylindole, dihydrochloride (DAPI) (ThermoFisher Scientific) for 5 minutes at RT. For mitochondrial staining, 5 × 104 live cells were incubated with the MitoTracker™ Red CMXRos FM (Invitrogen) in a glass bottom dish for 45 minutes. Cells were washed with x1 PBS 3 times and fixed with 4% formaldehyde for 15 minutes then washed by x1 PBS. Cells were incubated for 10 minutes in x1 PBS containing 0.1% Triton X-100 and stained with DAPI for 5 minutes at RT. Confocal images were acquired using a Zeiss LSM800 confocal laser scanning microscope (Carl Zeiss, Oberkochen, Germany). Confocal micrographs acquired with 10× and 20× objectives for phalloidin staining, and 40x for MitoTracker™ as a single optical slice.

### 17. Seahorse mitochondrial mito-stress test

Oxygen consumption rate (OCR) was measured using the Seahorse XF96 analyser (Agilent Technologies, Santa Clara, CA, USA) and Mito-stress test kit (Agilent Technologies) along with Seahorse XFe96/XF Pro FluxPak (Agilent Technologies) and Seahorse XF DMEM medium (without Phenol Red/pH 7.4/with HEPES/500 mL, Agilent Technologies). Initially, Mock^KD^ and YB^KD^ cells were seeded at 25,000 cells/well in 96-well plates on day one, while the XFe96 sensor cartridge hydrated in 200 µL calibration medium. On the next day, the cell medium was replaced with Seahorse XF DMEM, and cells were incubated at 37℃ in a CO₂-free incubator for 1 hour. During this time, 1 µM oligomycin, 0.5 µM FCCP, antimycin A, and rotenone were loaded into the drug ports of the cartridge. After loading, the sensor plate was calibrated in the analyser. Following calibration, the cell culture plate was loaded, and the analysis initiated with the program: set: Mixture 2 minutes, Measure 3 minutes, 4 sets in all steps. Ports: (A: Oligomycin / B: FCCP / C: Antimycin/Rotenone).

### 18. Puromycin incorporation assay

Nascent protein synthesis was evaluated by puromycin incorporation assay. Cells were treated with 10 μg/mL puromycin (Sigma-Aldrich) for 30 minutes prior to harvest. Cell lysates were prepared and analysed by western blot as previously described.

### 19. Statistical analysis

All experiments were done in 3 or more replicates. Statistical analysis was performed using GraphPad Prism 9.0. Analysis of two group comparisons was done using student’s t test and multiple group comparison was done using one-way ANOVA followed by Tukey’s post hoc test. p values of less than 0.05 were considered significant.

---

**Supplementary Fig 1:**
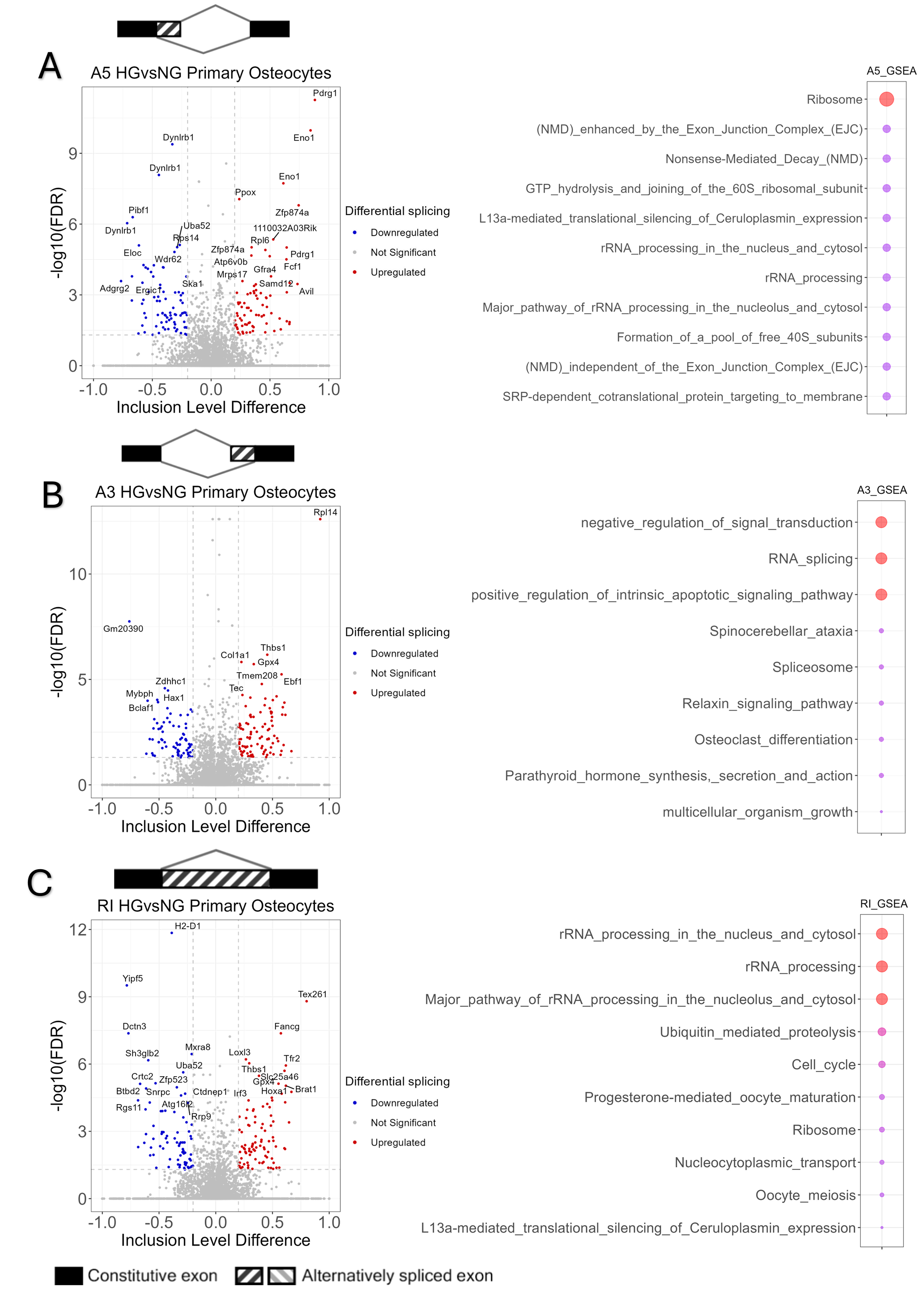
High glucose induces extensive AS changes in primary osteocytes. **(A-C)** Volcano plots and overrepresentation analysis of DSGs around AS events showing inclusion level (Δ1) difference comparing 5.5mM and 25mM glucose at FDR < 0.05 and 0.2 < Δ1 < -0.2 in primary osteocytes. Overrepresentation analysis was done through the eVITTA web-based toolbox. All analyses were done on n=3.

**Supplementary Fig 2:**
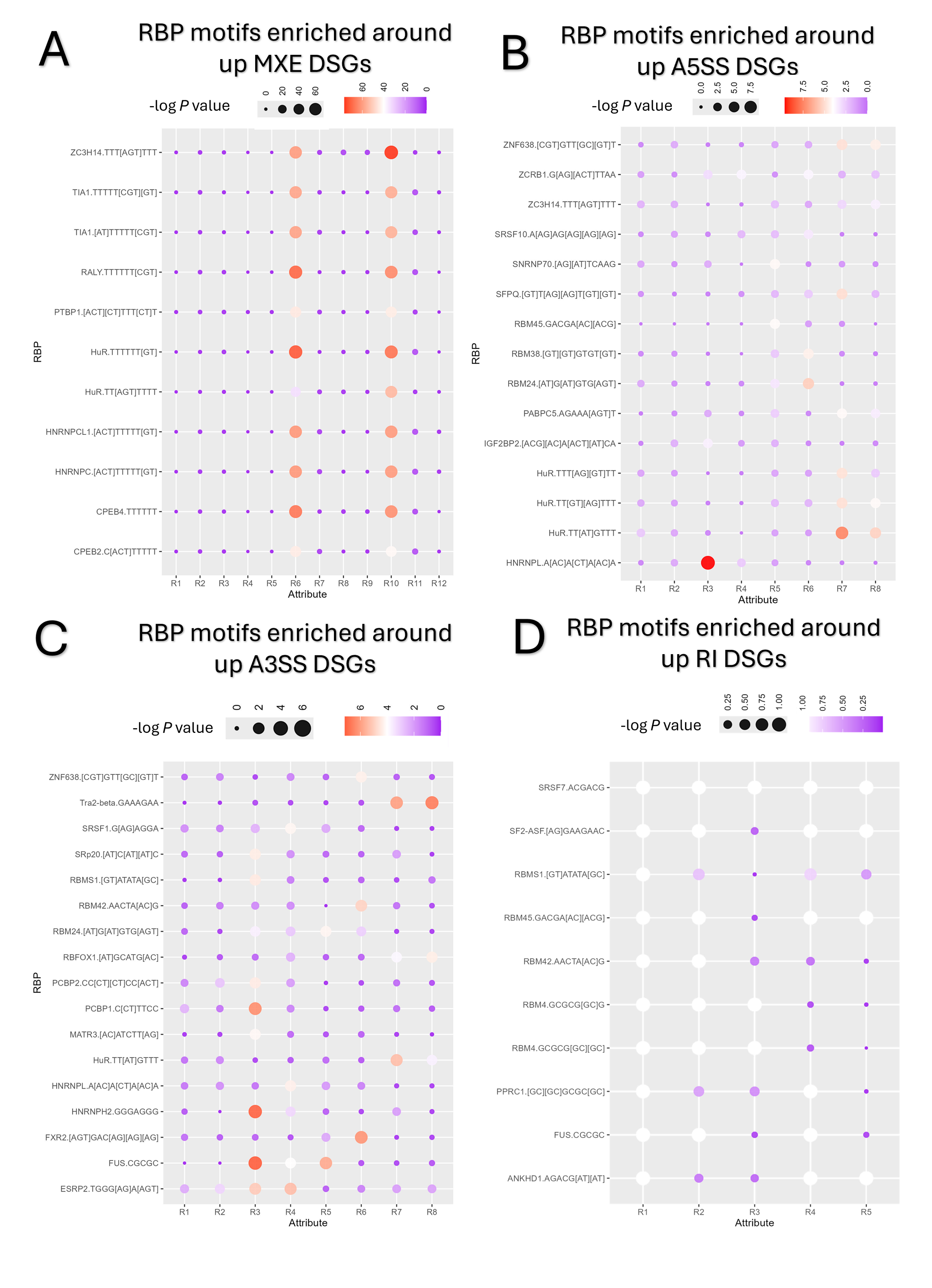
Significantly up-enriched RBPs motifs around AS events. **(A-D)** The x axis (Attribute) represents the position (R) of each splicing event in 5’ → 3’. RBP binding motifs are described in the y axis. The panels show MXE, A5SS, A3SS and RI RBP motifs around up and down regulated DSGs and bubble size and color denote the – log10 P value.

**Supplementary Fig 3:**
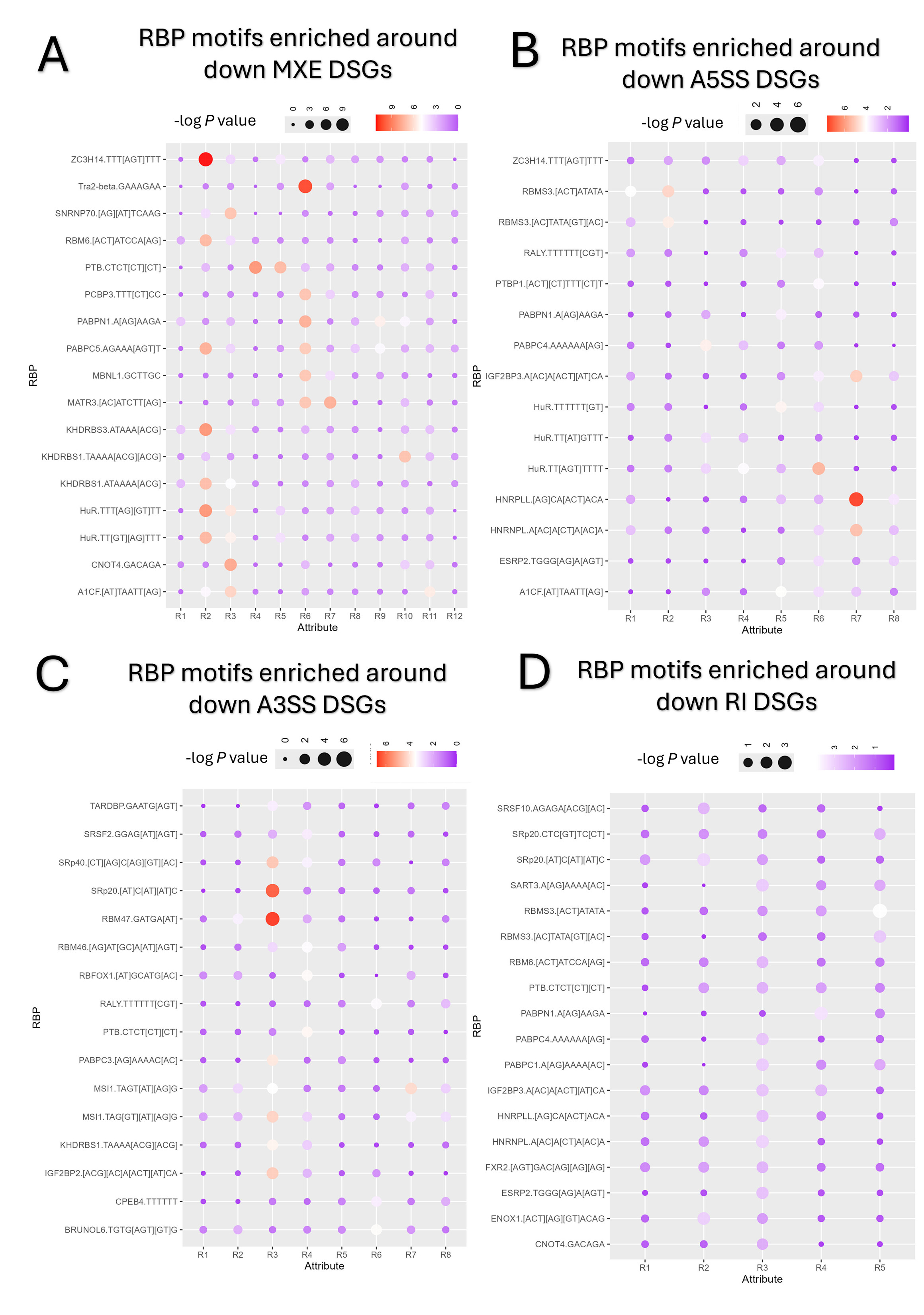
Significantly down-enriched RBPs motifs around AS events. **(A-D)** The x axis (Attribute) represents the position (R) of each splicing event in 5’ → 3’. RBP binding motifs are described in the y axis. The panels show MXE, A5SS, A3SS and RI RBP motifs around up and down regulated DSGs and bubble size and color denote the – log10 P value.

**Supplementary Fig 4:**
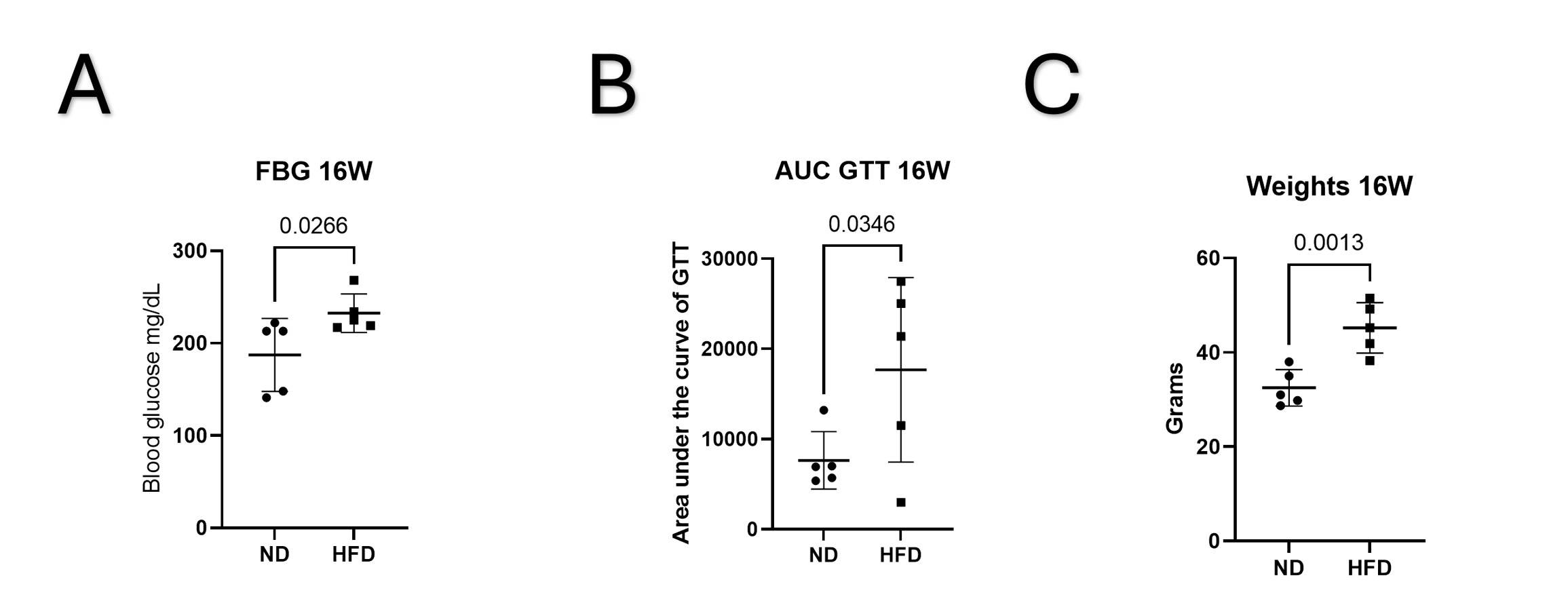
Weight and metabolic tests of mice fed a high fat diet for 16 weeks. **(A)** Six-hour fasted blood glucose levels of mice fed a ND or HFD. Data are presented as mean ± SD, n=5. **(B)** Glucose tolerance test area under the curve (AUC) calculated from individual glucose tolerance tests of mice fed a ND or HFD. Data are presented as mean ±SD, n=5. **(C)** Weight in grams of mice fed a ND or HFD Data are presented as mean ± SD, n=5.

**Supplementary Fig 5:**
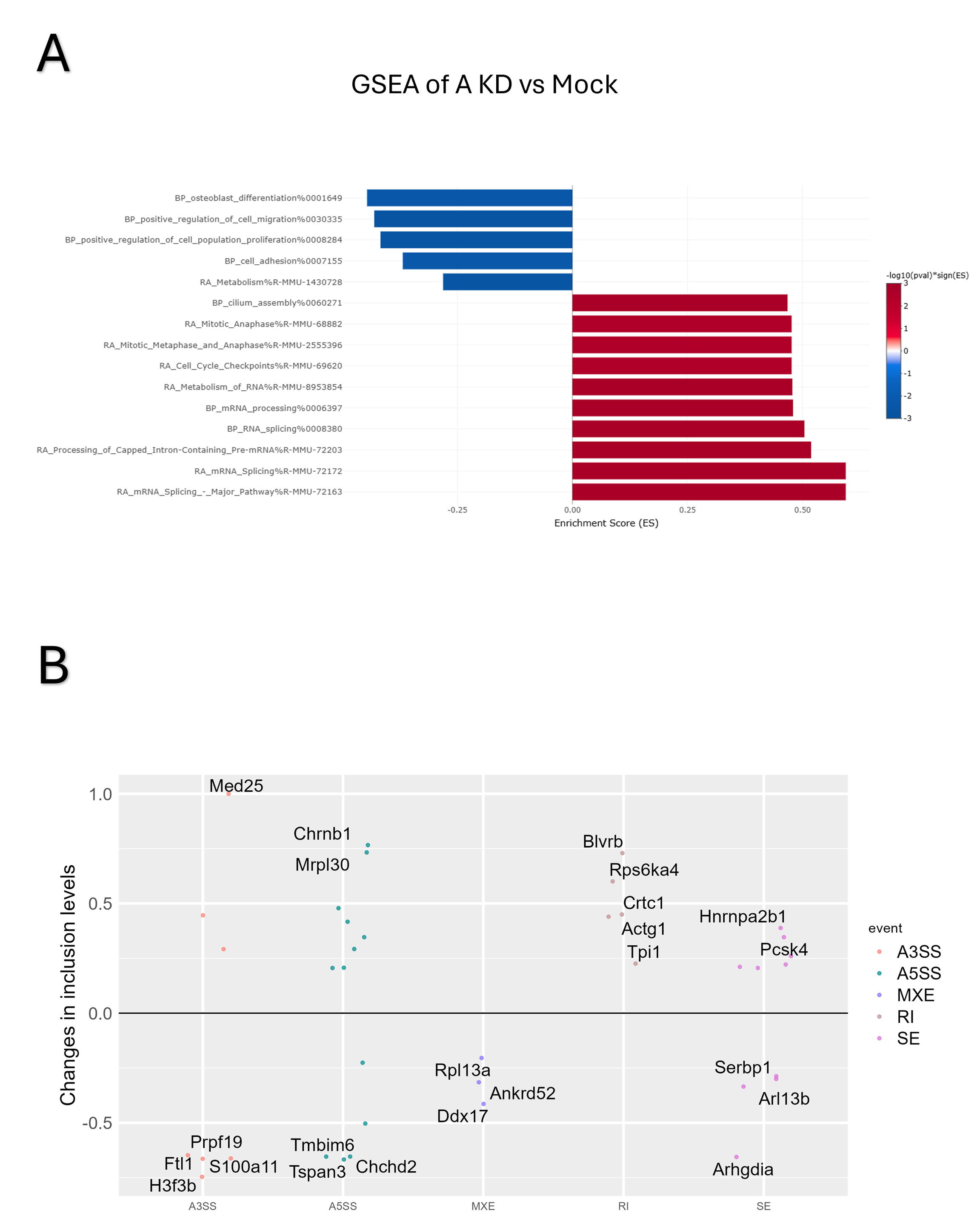
Alternative HuR KD (YA^KD^) construct enrichment analysis suggests off target effects. **(A)** Gene set enrichment analysis using eVITTA toolbox after YAKD of DEGs at LogFC 1 and -log10 adjusted p<0.05. Enriched terms include splicing and RNA metabolism as well as DNA damage repair, cell checkpoint activation and reduced cell adhension and proliferation. **(B)** Scatter plot showing DSGs around AS in YA^KD^ events in 5 patterns showing almost no differentially spliced genes at FDR < 0.05 and 0.2 < Δ1 < -0.2.

**Supplementary Table 1:** Over representation analysis of DSGs in primary osteocytes cultured in HG conditions.

**Supplementary Table 2:** Top 20 RNA binding motifs around differentially spliced transcripts in primary osteocytes cultured in HG conditions.

**Supplementary Table 3:** Over representation analysis of DEGS in after HuR KD.

**Supplementary Table 4:** DSGs, over representation analysis of DSGs and GOCC enrichment analysis after HuR KD.

**Supplementary Table 5:** Primers sequences, shRNA construct designs and antibodies used for Western blotting.

**Supplementary Table 6:** rMATS output from primary osteocytes cultured in HG conditions.

**Supplementary Table 7:** rMAPS output from primary osteocytes cultured in HG conditions.

**Supplementary Table 8:** rMATS output after HuR KD.

## Notes

**Funding statement:** This work is supported by The FRIS Creative Interdisciplinary Collaboration Program to AM and JST SPRING to ZF (No. JPMJSP2114).

https://dataview.ncbi.nlm.nih.gov/object/PRJNA1262104?reviewer=69lhvf8ud62ih53k7enfl6v3g0

### Competing Interest Statement

The authors have declared no competing interest.

### Summary of Updates

This version of the manuscript has been revised to update the following: Figure 6 with additional mechanistic data.

